# The Simultaneous Measurement of Reversed Phase-Encoding EPI in a Single fMRI Session: Evaluation of Geometric Distortion Correction in Submillimetre fMRI at 7T

**DOI:** 10.1101/2024.09.26.615120

**Authors:** Seong Dae Yun, Erhan Genc, Jeongbeen Lee, N. Jon Shah

## Abstract

The high temporal resolution of echo-planar imaging (EPI) has driven its widespread use in functional MRI (fMRI), and recent advancements in EPI have enabled the mapping of cortical layer-specific functional activities. Notwithstanding these technical innovations, geometric distortion correction in EPI remains essential to ensure accurate functional mapping onto anatomical references. A common approach for distortion correction involves the acquisition of an additional scan, with the phase-encoding direction reversed. However, this extra scan necessitates redundant acquisitions of calibration scans, significantly increasing acquisition time and energy deposition. To address this concern, this work presents a novel EPI scheme that simultaneously acquires both the original and reversed phase-encoding data within a single fMRI session. Furthermore, despite the widespread use of distortion correction, methods for qualitative and quantitative evaluation of its impact on submillimetre fMRI analysis have remained largely unexplored. This study acquires submillimetre visual fMRI data (0.73 × 0.73 mm^2^) at 7T using the suggested EPI scheme and evaluates the impact of distortion correction through various metrics proposed here, including spatial resolution, co-registration accuracy, functional mapping fidelity, and distribution of functional voxels. Our fMRI acquisition scheme effectively reduced redundant acquisition time and total energy deposition to subjects. Distortion-corrected fMRI data demonstrated significant improvements in co-registration with anatomical scans, thereby enhancing functional mapping accuracy. The improvements were achieved without significant degradation of spatial resolution or alteration of the functional activation distribution. These findings were verified through qualitative and quantitative assessments, highlighting the effectiveness of distortion correction in submillimetre fMRI.

**Highlights:**

- We present a novel fMRI scheme that effectively acquires reversed-PE EPI data
- Our fMRI scheme substantially reduces extra acquisition time and energy deposition
- The impact of EPI distortion correction is evaluated for submillimetre fMRI
- We propose diverse qualitative and quantitative metrics, enabling thorough analysis
- Distortion correction demonstrates its effectiveness for high-fidelity fMRI mapping

## 1. Introduction

The advent of echo-planar imaging (EPI) (Mansfield, 1977) has facilitated the development of various dynamic MRI applications. Its relatively high temporal resolution makes it particularly well-suited for detecting time-dependent variation in signal contrast, leading to its widespread use in applications such as diffusion and functional MRI (fMRI) studies. Recent advancements in EPI techniques and associated image analysis methods have now enabled the characterisation of physiological quantities in the brain with respect to the cortical depths, providing deeper insights into brain function and clinical abnormalities.

Despite these technical innovations, the relatively low receiver bandwidth along the phase-encoding (PE) direction in EPI significantly contributes to the development of geometric deformation of tissue structures, making image analysis based on accurate anatomical references challenging. This geometric distortion can be effectively mitigated through technical approaches such as improved shimming (Balteau et al., 2010; Kim et al., 2016) or readout-acceleration techniques, including parallel imaging, partial Fourier, or multi-shot methods. However, they may necessitate technically more demanding configurations, especially when operating at higher field strengths or aiming for higher-resolution imaging with an increased readout length.

Alternatively, a widely adopted methodological approach to address the EPI distortion is the direct unwarping of geometric distortions using an additional EPI scan acquired with the reversed PE direction (Chang and Fitzpatrick, 1992; Morgan et al., 2004), commonly referred to as the ‘blip-up-blip-down’ method. This approach exploits the fact that distortions in the reversed PE data develop toward the opposite direction, enabling the data-driven calculation and relocation of voxel shifts between the two different PE directions without prior knowledge of the B_0_ offset (Jezzard and Balaban, 1995; Chen and Wyrwicz, 1999; Hutton et al., 2002) or PSF (Robson et al., 1997; Zeng and Constable 2002; Zaitsev et al., 2004; Chung et al., 2011; Patzig et al., 2021). Moreover, the reversed PE data can be obtained using the same sequence, indicating that no additional sequence implementation is required, unlike the B_0_ and PSF methods. Several comparative studies have demonstrated the superior performance of the blip-up-blip-down method over the B_0_ field map method (Wang et al., 2017; Malekian et al., 2023).

However, one concern that has not been extensively explored in the blip-up-blip-down method is the potential increase in redundant acquisition time and specific absorption ratio (SAR) accrued during the acquisition of reversed-PE EPI. This arises due to the fact that the reversed PE data are typically acquired by adding an extra run of the same EPI protocol, only with the change in the PE direction, resulting in a repeated measurement of calibration scans for image reconstruction. To address this issue, this work presents a novel EPI scheme that additionally acquires EPI data with the reversed PE direction within a single fMRI session, thereby facilitating the analysis of distortion-corrected images with reduced motion effects. This proposed scheme is expected to shorten the total acquisition time while reducing energy deposition, particularly for fMRI, which requires a large imaging matrix size to achieve high spatial resolution and whole-brain coverage.

In addition, notwithstanding the widespread use of the blip-up-blip-down method for the analysis of submillimetre fMRI data (Kashyap et al., 2018; Marquardt et al., 2018; Jia et al., 2020; Huang et al., 2021; Navarro et al., 2021; Schellekens et al., 2023; Yun et al., 2023), the evaluation of the distortion-correction performance and its potential impact on fMRI data has remained largely unexplored in the context of the submillimetre fMRI. While several previous studies have demonstrated the effectiveness of distortion correction (Hutton et al., 2002; Dymerska et al., 2018; Duong et al., 2020; Patzig et al., 2021; Montez et al., 2023), most have primarily focused on structural improvements in distortion-corrected EPI and its co-registration accuracy with anatomical scans, often overlooking the impact of distortion correction on high-resolution functional data.

Another objective of this work is to propose quantitative and qualitative metrics for the comprehensive evaluation of the performance of distortion-corrected fMRI data in terms of structural and functional mapping quality. This includes the assessment of potential spatial resolution degradation, which is crucial for submillimetre fMRI analysis, the examination of structural alignment between functional and anatomical scans, and the evaluation of functional mapping accuracy along with a statistical comparison of activated voxels. This study validates the feasibility of the proposed fMRI scheme and the performance evaluation metrics using submillimetre (0.73 × 0.73 mm^2^) fMRI data acquired on a 7T MRI scanner.

## 2. Materials and Methods

### 2.1. Study subject and ethics statement

All in vivo measurements were performed on healthy volunteers who underwent standard safety screening procedures. None had any medical conditions or suffered from any neurological or psychiatric illnesses. After receiving a complete explanation of the study, all participants provided written informed consent before scanning. The study protocol, screening questionnaires, and consent forms were approved by the local institutional review board at RWTH Aachen University, Germany (EK 346/17). Five healthy volunteers participated in this study, during which visual stimuli were applied.

### 2.2. Proposed EPI scheme

Figure 1a shows a schematic representation of a typical fMRI run using the standard EPI scheme, consisting of three stages: calibration, dummy, and actual fMRI scans. Before starting the actual fMRI scans, two preceding measurements, namely calibration and dummy scans, are performed, which are necessary for the reconstruction of acceleration techniques and for reaching a steady state (Yun et al., 2023). The acquisition of calibration data is assumed to be included here, as most high-resolution fMRI protocols commonly employ acceleration techniques such as parallel imaging. In the standard scheme, EPI with reversed PE is typically performed by adding a second run of the same protocol, differing only in the PE direction; see the blue and red arrows indicating the PE directions in the fMRI and reversed PE measurements, respectively. This approach suggests that the second session still requires another complete acquisition of the calibration data, although the reversed-PE EPI data only need to be reconstructed for a few temporal volumes.

**Figure 1.**
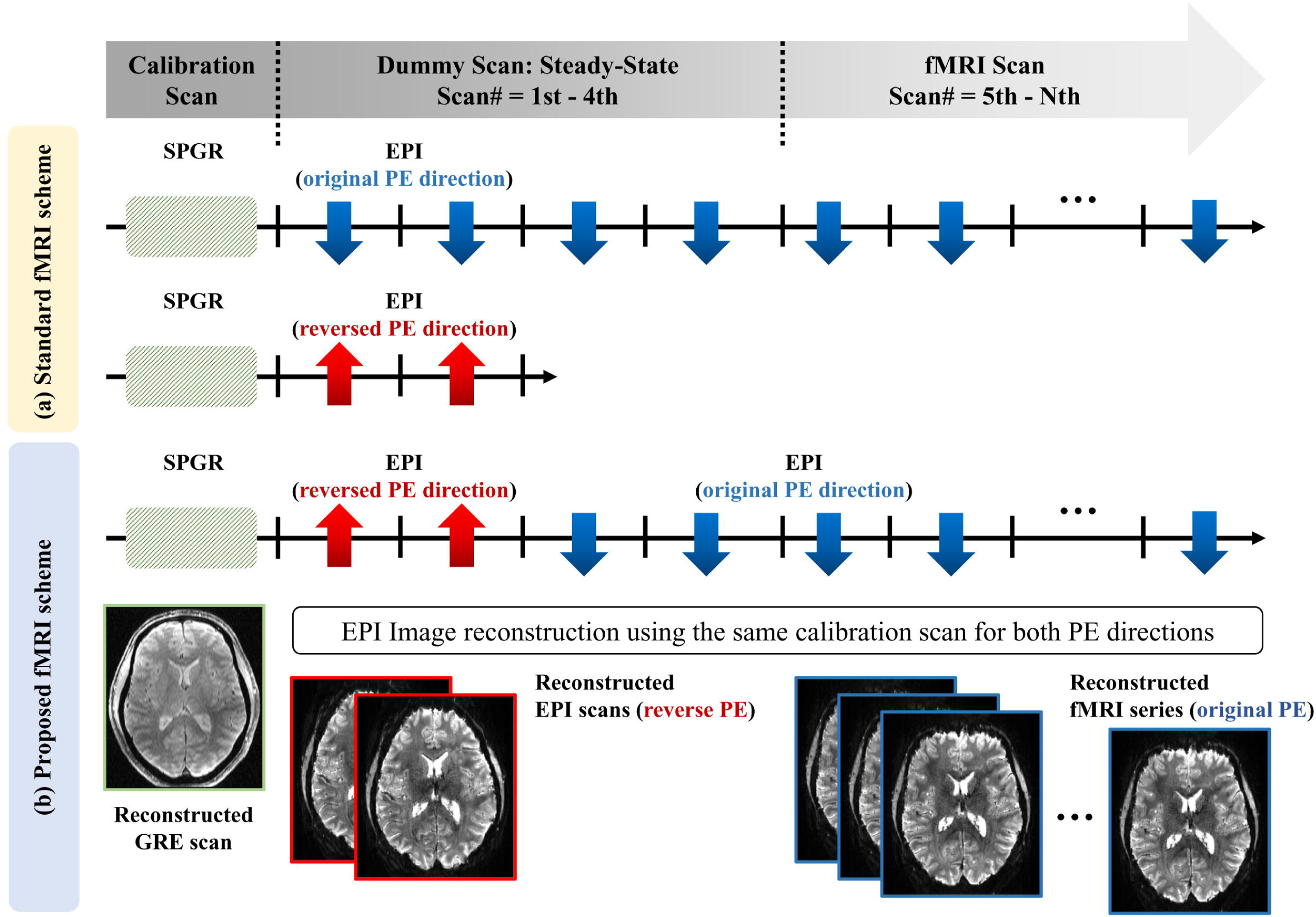
Schematic diagram of an fMRI acquisition sequence. (**a**) Standard EPI and (**b**) proposed EPI. In the standard scheme, the reversed PE data are acquired in a separate session, requiring repeated measurements for the calibration scan. In contrast, the proposed scheme acquires the reversed PE data during the same session at the dummy stage, allowing the same calibration scan to be used for image reconstruction of both original and reversed-PE data. The blue and red arrows denote the original and reversed PE directions, respectively. In this work, the number of dummy and reversed PE scans was configured to four and two, as illustrated in the figure.

While the fMRI scan is acquired using EPI, the calibration scan for the parallel imaging is often acquired using sequences more tolerant to field inhomogeneities, such as a conventional line-by-line excitation, spoiled gradient-recalled (SPGR) sequence, ensuring more robust kernel calibration at ultra-high fields. However, the SPGR scheme requires a relatively long acquisition time, making the repeated acquisition of the calibration scan in the standard scheme increasingly redundant. To resolve this issue, the proposed scheme acquires the first two dummy scans– excluded from image analysis– using reversed PE, while the remaining scans are acquired with the original PE directions (Yun et al., 2023). This strategy allows a single calibration kernel to be shared to reconstruct both the original and reversed-PE EPI data, effectively reducing the total acquisition time and SAR for parallel imaging calibration. In this work, the number of dummy and reversed PE scans was set to four and two, respectively, as illustrated in the figure.

### 2.3. Imaging protocols: submillimetre visual fMRI

A visual checkerboard paradigm was employed for in vivo fMRI experiments to elicit circumscribed activation in the visual cortex (Yun et al., 2013, 2017). The passive viewing task involved a simple block design in which a black and white checkerboard, reversing at a frequency of 8 Hz, alternated with low-level baseline phases (fixation cross). The paradigm consisted of four dummy scans for reaching a steady state, followed by 96 scans comprising eight cycles of baseline-activation states. Each cycle lasted six volume acquisitions, resulting in a total duration of 21 seconds. As described in the previous section, the first two dummy scans were acquired with the reversed PE direction.

The feasibility of using the proposed EPI scheme for visual fMRI was verified with the following imaging parameters: TR/TE = 3500/22 ms, FOV (AP × LR × FH) = 210lJ×lJ210 × 117lJmm3, matrix = 288lJ×lJ288lJwithlJ117 axial slices (0.73lJ×lJ0.73lJ×lJ1.0lJmm3), flip angle = 90°, partial Fourier = 5/8, parallel imaging = 3-fold, multi-band = three-fold, and receiver bandwidth = 965 Hz/Px. The parallel imaging calibration data were acquired using the SPGR sequence with an auto-calibration signal (ACS) size of 72. For the assessment of reversed PE data from the proposed scheme, additional reversed PE data were separately acquired using the standard scheme (see Fig. 1b). A 3D anatomical image was acquired using a T1-weighted magnetisation-prepared, rapid gradient echo (MP2RAGE) sequence with the following configuration: TR/TE/TI1/TI2 = 4440/2.08/840/2370 ms, FOV (FH × AP × LR) = 240lJ×lJ226 × 154lJmm3, matrix size = 400lJ×lJ376 with 256 sagittal slices (0.6 mm isotropic), flip angles = 5°/6°, parallel imaging = 3-fold, and receiver bandwidth = 250 Hz/Px. The above imaging setup was implemented on a Siemens Magnetom Terra 7T scanner with a single-channel Tx and 32-channel Rx Nova medical coil supplied by the manufacturer.

### 2.4. Distortion correction: blip-up-blip-down method

Geometric distortion in the original fMRI time-series data (96 temporal volumes) was corrected using reversed-PE data (two temporal volumes in the dummy can) acquired from the proposed scheme. This blip-up-blip-down method was implemented using a publicly available image processing toolkit, Advanced Normalization Tools (ANTs) (https://github.com/srikash; https://github.com/ANTsX/ANTs; Avants et al., 2010; Esteban et al., 2019; Tustison et al., 2021). ANTs computes the warping field of the original EPI data while simultaneously estimating subject motion displacements, enabling correction of both motion and distortion within the time series in a single interpolation step. This strategy assists in minimising potential spatial resolution degradation, which may be more pronounced when motion and distortion are corrected separately (Glasser et al., 2013; Wang et al., 2022).

To validate the use of reversed-PE data from the proposed scheme, the same ANTs correction process was repeated using reversed-PE data acquired with the standard scheme in a separate session (see Fig. 1a). Differences in motion displacement between the proposed and standard schemes were examined, and their impact on distortion correction was assessed.

### 2.5. Performance evaluation: qualitative and quantitative metrics

#### 2.5.1. Spatial resolution

It has been previously reported that the distortion correction step involves resampling of voxels or sub-voxel shifts, which can result in a loss of effective spatial resolution (Renvall et al., 2016; Kashyap et al., 2018; Polimeni et el., 2018; Wang et al., 2022). This study thoroughly investigates the potential degradation of spatial resolution caused by the unwarping process during distortion correction, building upon and modifying an approach proposed in previous work (Renvall et al., 2016).

The original EPI image was subjected to the standard spatial smoothing procedure in SPM12 (Wellcome Department of Imaging Neuroscience, UCL, London, UK; https://www.fil.ion.ucl.ac.uk/spm/software/spm12/) using Gaussian smoothing kernel widths (σ) ranging from 0.2 to 0.8 mm in 0.1 mm increment. A region-of-interest (ROI) was selected in the white matter (WM), where the signal variation in the region was relatively homogeneous. The standard deviation (SD) within the ROI was computed for the original and the smoothed EPI images, creating a lookup table to examine the spatial resolution based on the SD values. A lower SD indicated greater spatial resolution degradation due to more intensive Gaussian smoothing with a larger kernel size. The SD was then computed for the distortion-corrected EPI image and compared to the lookup table to identify the closest match. The corresponding smoothing kernel size was used to estimate the degree of the spatial resolution degradation (σ_deg_) in the distortion-corrected image, as defined by the following formula:

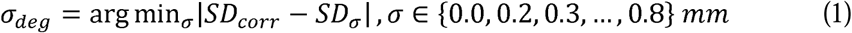

where *SD_corr_* and *SD*_σ_ represent the SD from the distortion-corrected image and the original EPI images smoothed with different kernel widths, respectively, with σ = 0.0 indicating no smoothing. σ*_deg_* denotes the kernel width creating the closest match. For each fMRI data set, this computation was performed for every slice location, and the results were averaged across slices.

#### 2.5.2. Structural similarity: co-registration

The original and distortion-corrected EPI images were independently co-registered to the MP2RAGE scan using the ‘antsRegistrationSyN’ module in the ANTs toolkit. In each case, the structural similarity between the EPI and the MP2RAGE scans was visually inspected by comparing the co-registered anatomical structures, particularly in regions like the frontal lobe where EPI distortion is most pronounced. The similarity was further quantified using the Learned Perceptual Image Patch Similarity (LPIPS) index, a perceptual metric that measures image similarity by computing deep feature distances across multiple layers of a pre-trained neural network (Zhang et al., 2018). LPIPS captures perceptual differences by comparing feature maps extracted from various convolutional layers and aggregating their distances. Compared to traditional similarity metrics, such as L2, Structural Similarity Index Measure (SSIM) (Wang et al., 2004), and Feature Similarity Index Measure (FSIM) (Zhang et al., 2011), LPIPS exhibits a significantly stronger correlation with human perceptual judgments, making it a more robust evaluation metric for image similarity. In this work, LPIPS was configured with the Visual Geometry Group (VGG) network, which has been shown to outperform shallower, more lightweight architectures like SqueezeNet in terms of perceptual similarity (Zhang et al., 2018).

For this comparison, the standard segmentation routine in SPM12 was applied individually to the EPI and MP2RAGE scans, generating probability maps for grey matter (GM), WM, and cerebrospinal fluid (CSF) masks. The probability maps obtained were then used to compute the LPIPS index as shown in the formula below, where a lower value indicates higher image similarity:

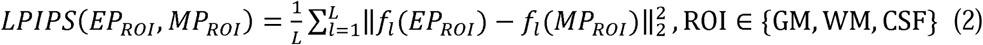

where *EP_ROI_* and *MP_ROI_* represent the probability maps of EPI and MP2RAGE at the selected ROI, respectively. *f_l_*denotes the deep feature map extracted from the *l*-th convolutional layer, and *L* is the total number of layers considered in the computation. The computation was applied for each slice, and the average score was obtained. This approach allows for the comparison of structural similarity between the EPI and MPRAGE scans, each having different contrasts (i.e. T_2_* and T_1_). Prior to data processing, background noise in the MP2RAGE scan, which was generated by combining the two inversion images into a single unified image, was removed using an open-source script (https://github.com/JosePMarques/MP2RAGE-related-scripts).

#### 2.5.3. Functional mapping

For the original and distortion-corrected fMRI data sets, the first-level analysis was performed separately using the generalised linear model in SPM12, and the functional results obtained with an uncorrected *p*-value < 0.001 were overlaid on their respectively co-registered MP2RAGE scan for visual inspection. From the total number of activated voxels, the proportion located within the GM, as defined by the MP2RAGE scan, was calculated. To

minimise false positives in this analysis, only activated voxels with t-values of five or greater were included (Lohmann et al., 2018):

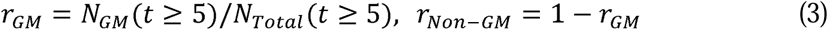

where *r_GM_* and *r_Non-GM_* represent the ratio of activated voxels within the GM and non-GM regions, respectively. The ratios were computed for the results obtained from both the original and distortion-corrected fMRI datasets, and putative differences were examined.

Lastly, the impact of the distortion correction on the functional activation was inspected by analysing the histogram distribution of the *t*-values for the activated voxels. The histogram results obtained from the original and distortion-corrected fMRI data were compared, and the degree of matching between the two cases was quantitatively measured by computing the Pearson correlation coefficient:

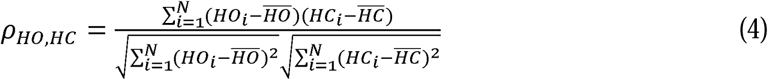

where *HO_i_* and *HC_i_* represent the values of the *i*-th bin in the original and distortion-corrected histograms, respectively. *N* denotes the number of bins in the histograms. The differences in t-value distribution before and after distortion correction were statistically assessed using the Kolmogorov-Smirnov (KS) test:

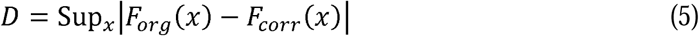

where *F_org_(x)* and *F_corr_(x)* represent the cumulative distribution function of the t-values from the original and distortion-corrected results. *D* is the KS statistic that reflects the degree of discrepancy between the two distributions, with values ranging from 0 to 1. The corresponding *p*-value was also computed.

## 3. Results

### 3.1. Distortion correction using the proposed scheme

Figure 2 shows the results of distortion correction for fMRI scans at four representative slice locations, with the first and second rows displaying images obtained from the reversed (1st scan) and original PE (5th scan) directions. The last row shows the corresponding distortion-corrected results, revealing no significant visible loss of spatial resolution at any of the displaced slice locations. This result also demonstrates the feasibility of using the proposed fMRI scheme in the distortion correction. The distortion-corrected results at entire slice locations can be found in the Supplementary File (see Supplementary Figure 1).

**Figure 2.**
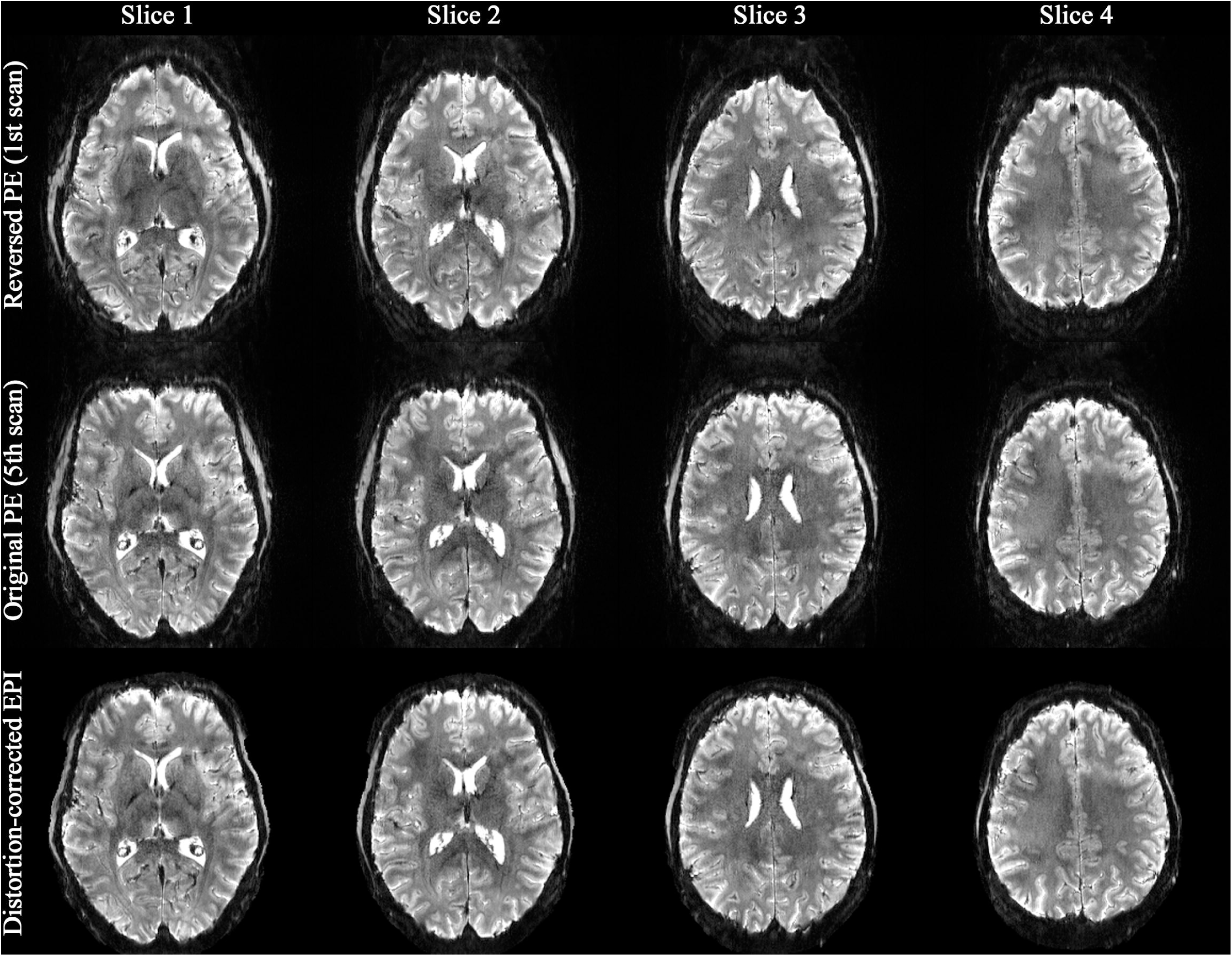
Reconstructed images at four representative slice locations. The first and second rows display images obtained from the reversed and original PE directions. The last row shows the corresponding distortion-corrected results, revealing no significant visible loss of spatial resolution.

Figure 3a displays reversed PE images obtained from both the standard and proposed schemes at a representative slice location. Figure 3b highlights the ventricle boundaries extracted from the reversed-PE images from the standard (yellow), overlaid on those from the proposed (magenta) schemes. This image panel illustrates the significant motion displacement that occurred between the fMRI session and the subsequent session for acquiring reversed-PE data in the standard scheme. The corresponding distortion-corrected results, obtained using both reversed-PE images, are shown in Fig. 3c. Both methods effectively eliminate the geometric distortions present in the original EPI image (Fig. 3d). However, the proposed method demonstrates greater improvement in signal recovery around the frontal lobe, as well as the clearer delineation of the GM and WM boundaries (see the regions marked by arrows in Fig. 3e). Furthermore, the proposed simultaneous acquisition scheme significantly reduces the redundant SAR and acquisition time for the acquisition of reversed-PE EPI. A more detailed analysis, including the technical aspects of the imaging sequences, is presented in the supplementary Table 1.

**Figure 3.**
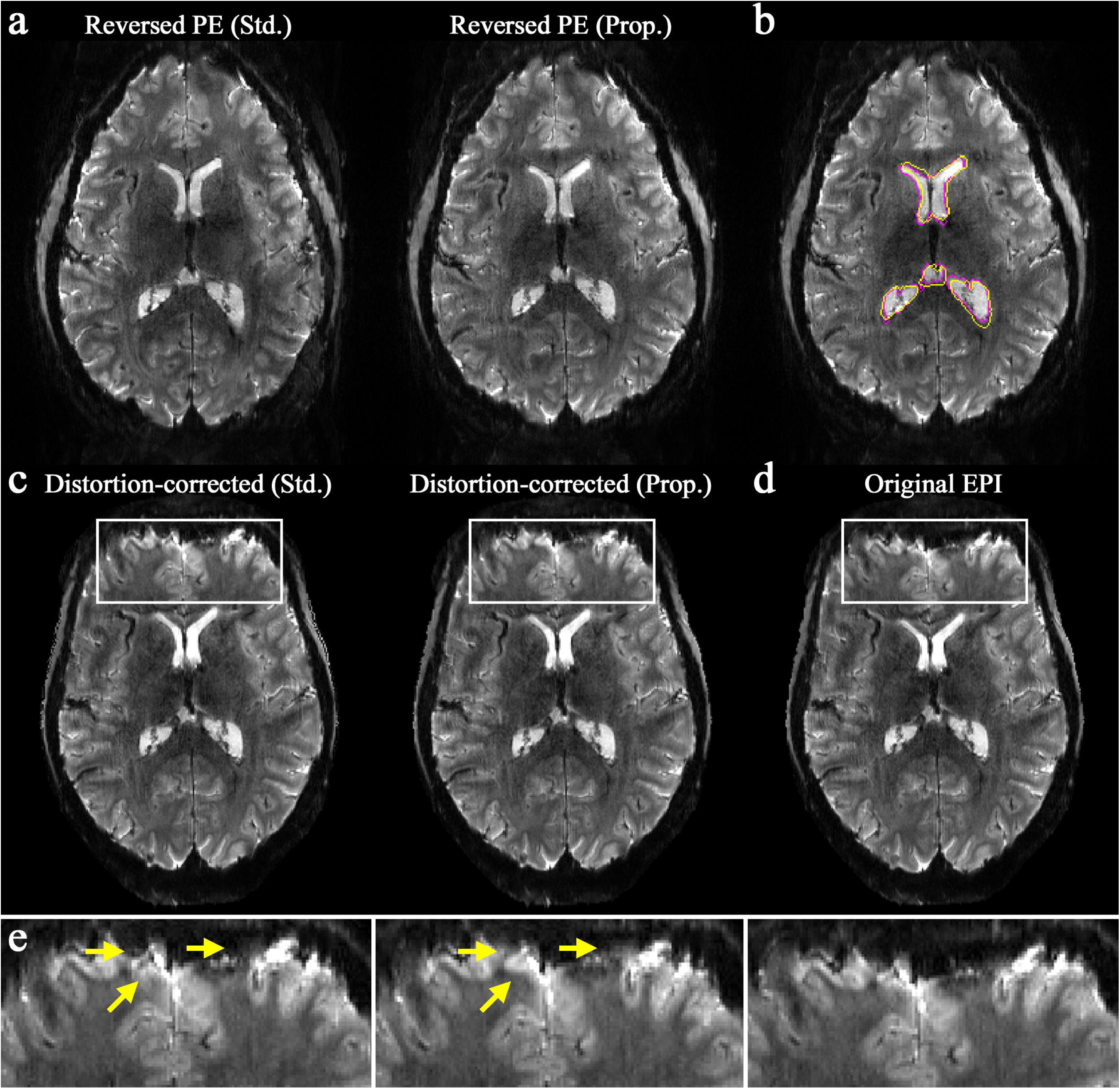
Effects of subject motion on distortion correction. (**a**) Reversed PE images obtained using the standard (left) and proposed (right) schemes. (**b**) Ventricle boundaries extracted from both images in Fig. a, shown in yellow (standard) and magenta (proposed), overlaid on the reversed PE image from the proposed method, illustrating significant motion displacement between the two cases. (**c**) Corresponding distortion-corrected images generated from the reversed PE images in Fig. a. (**d**) Original, uncorrected EPI image. (**e**) Enlarged depiction of the rectangular ROIs, highlighting effective distortion correction for both the standard and proposed schemes. However, the proposed scheme demonstrates greater improvement in signal recovery and delineation of GM and WM boundaries around the frontal lobe (see arrows).

### 3.2. Assessment of spatial resolution

Figure 4a displays the original and distortion-corrected EPI images at a representative slice location. The yellow ROIs indicate the region used for the SD computation in the quantitative assessment of spatial resolution. The rectangular ROIs mark the areas where spatial resolution was visually inspected, with an enlarged view provided in Fig. 4b. The enlarged views of the original image, outlined in white rectangular ROIs, illustrate the effects of various smoothing kernel widths, where a decrease in spatial resolution becomes more apparent at larger kernel widths. For the selected slice location, visual inspection of Fig. 4b suggests that the loss of spatial resolution induced by distortion correction was not significant.

**Figure 4.**
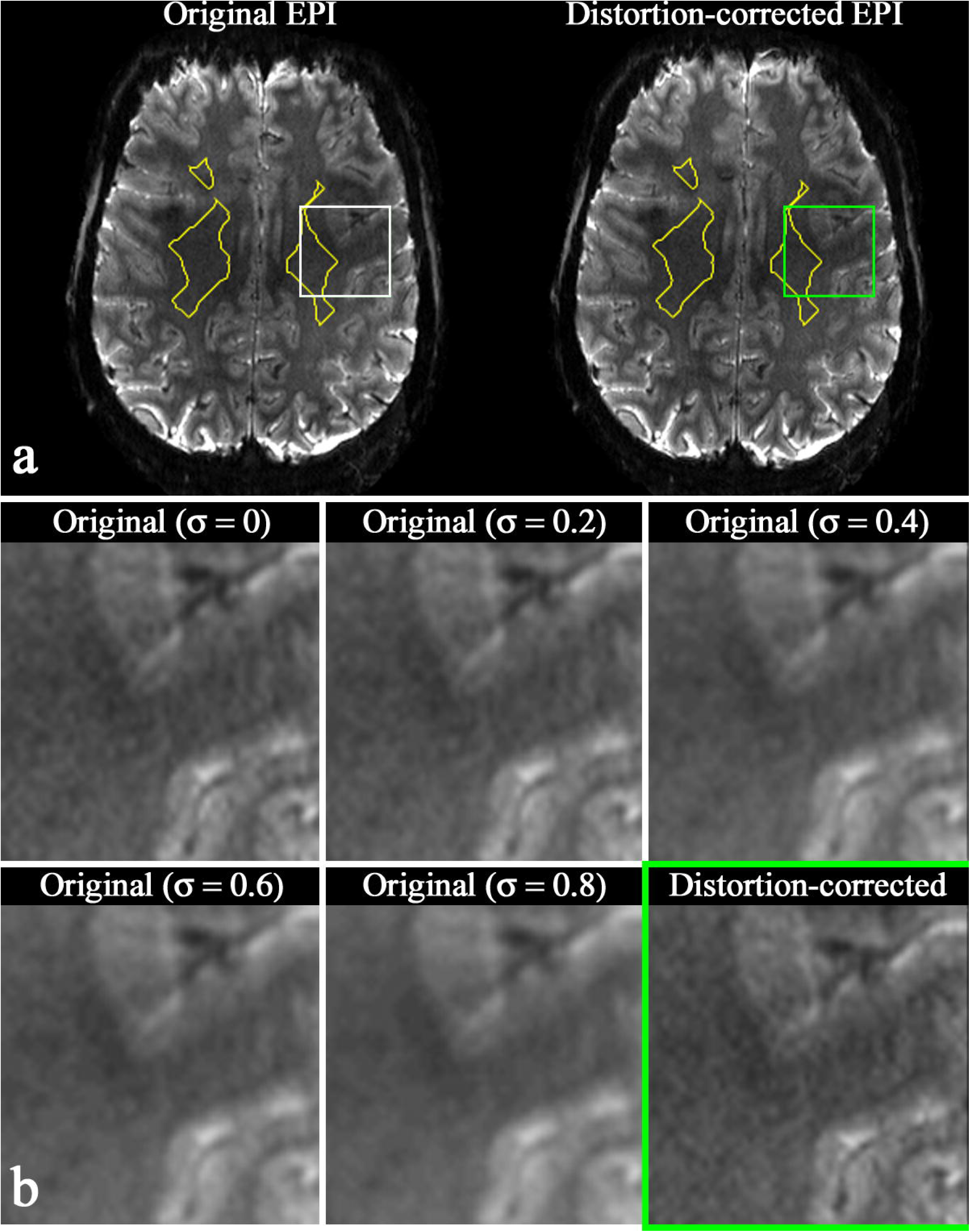
Results of spatial resolution assessments. (**a**) Original and distortion-corrected EPI images at a representative slice with selected ROIs: yellow for SD computation, and white (original) and green (distortion-corrected) for visual inspection. (**b**) From top-left to bottom-right, enlarged depictions of the rectangular ROIs are shown for the following: original EPI, original EPI smoothed with Gaussian kernel widths (σ) of 0.2, 0.4, 0.6 and 0.8, and distortion-corrected EPI. Visual inspection of image panel b suggests that the loss of spatial resolution in the distortion-corrected image is not significant.

Table 1 presents the obtained lookup table at the above slice location, listing the SD values of the original image and the spatially smoothed ones. The SD value of the corresponding distortion-corrected image was computed as 72.55, which is closest to that of the original image without smoothing. Consequently, the degree of the spatial resolution degradation (σ_deg_) was determined to be 0.0. The results of σ_deg_ obtained at all slice locations are presented in the box-plot in Figure 5. The results from four out of five subjects show that the median value was zero, with the third quartile ranging between 0.2 and 0.3. While the third subject showed a relatively higher third quartile, the median value remained low at around 0.3. These findings suggest that the spatial resolution degradation caused by distortion correction was not significant.

**Figure 5.**
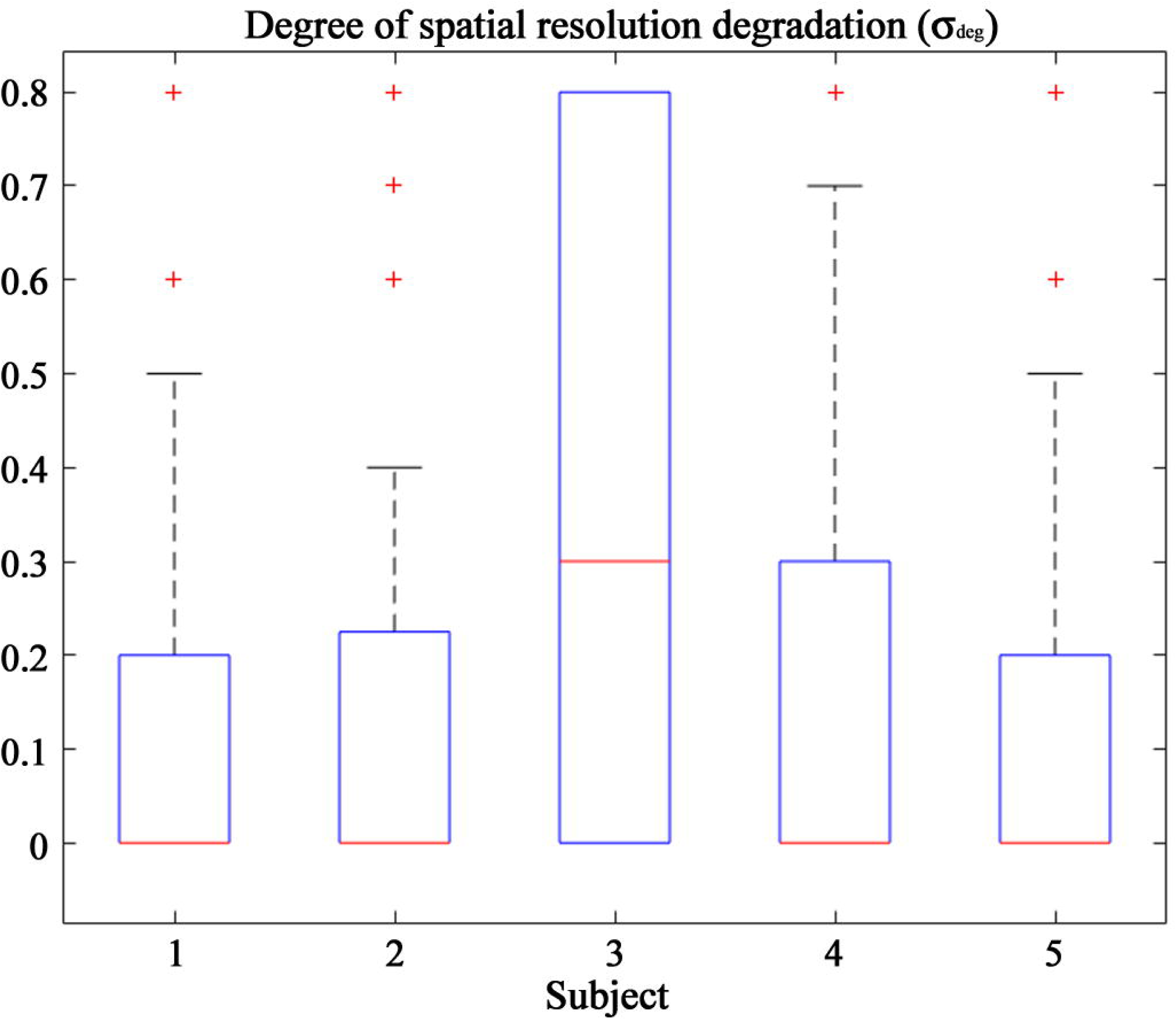
Box-plot of the spatial resolution degradation (σ_deg_) across five subjects. The results indicate that the spatial resolution degradation caused by distortion correction was not significant.

**Table 1.**
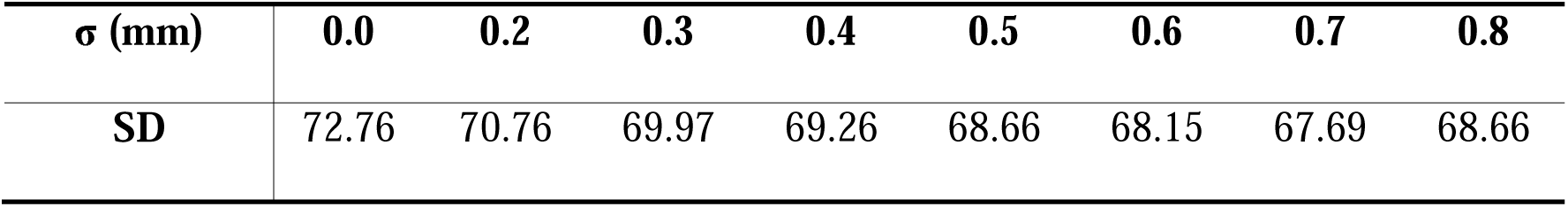
Lookup table showing the SD values of the original image, smoothed with a range of kernel widths (σ), at the specified slice location (see Fig. 4). The SD of distortion-correction image was computed as 72.55, which is closest to the original image with no smoothing applied (i.e. first column).

### 3.3. Co-registration between the functional and anatomical scan

The EPI images, co-registered with the MP2RAGE scan, were transformed from the subject space to the anatomical space, and results from a representative subject are shown in Fig. 6. Both the EPI and MP2RAGE scans are presented in three sectional views (i.e. axial, coronal and sagittal), each depicting the counterpart of the brain. That is, for the axial and coronal presentation, the EPI and MP2RAGE scans display the left and right hemispheres, whereas they show the upper and lower halves for the sagittal presentation, respectively.

**Figure 6.**
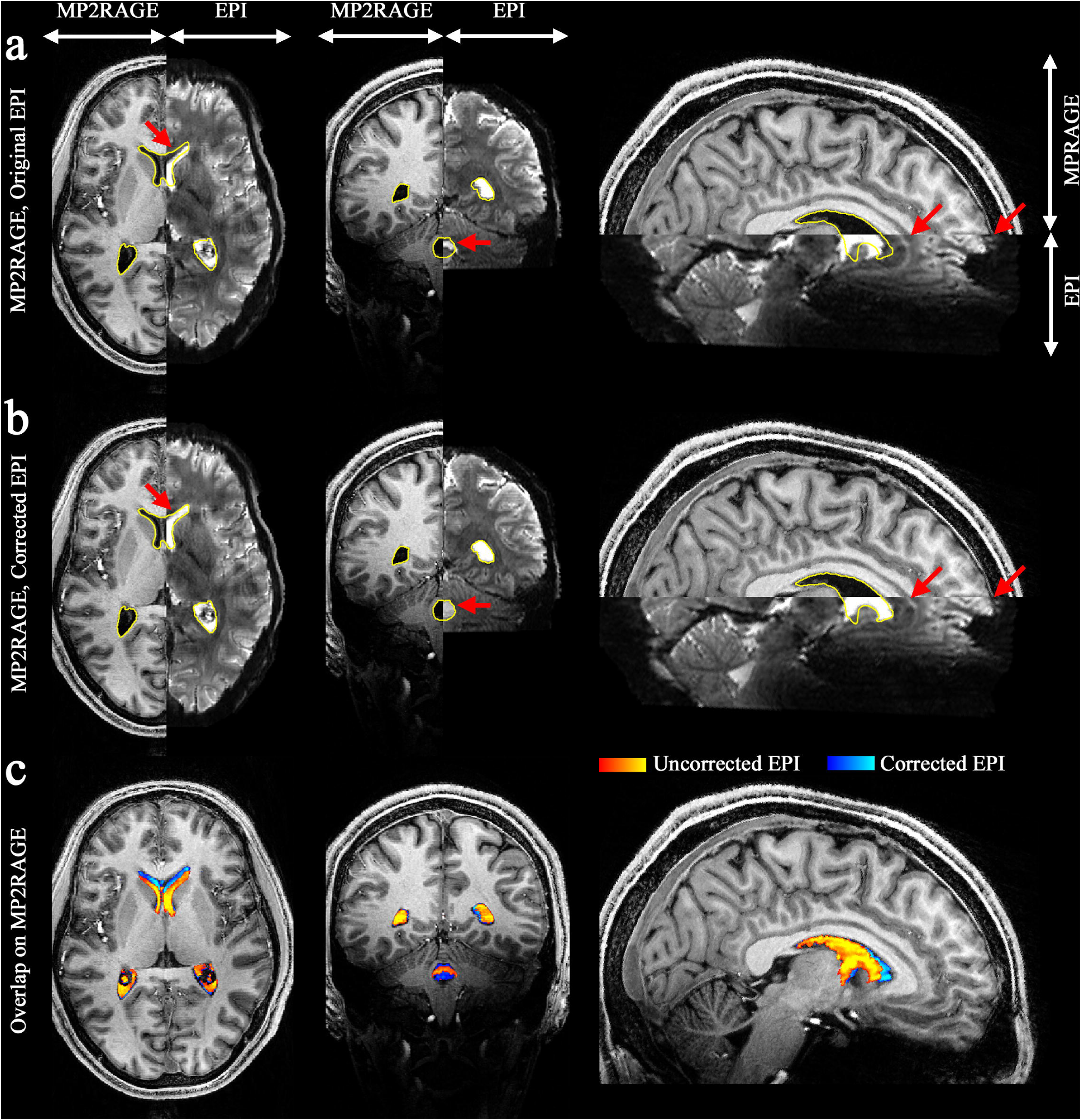
Results of co-registration to the anatomical scan (MP2RAGE). (**a**) Original EPI, (**b**) distortion-corrected EPI and (**c**) both results overlaid onto the MP2RAGE. In image panels a and b, the EPI and MP2RAGE scans are displayed alongside, each depicting the counterpart of the brain. The results from distortion-corrected EPI show improved alignment with the MP2RAGE scan, particularly around the regions marked by yellow boundaries (i.e. lateral and fourth ventricles) and red arrows; the boundary between the cingulate gyrus and corpus callosum is also shown to be well aligned. The overlapped depiction in image panel c further confirms the improved co-registration, where the ventricle signals are represented in red-yellow for the original EPI and blue-light blue for the corrected EPI.

To aid visual comparison, the contour of the ventricles (i.e. the lateral and fourth ventricles) obtained from the MP2RAGE scan are depicted in yellow. Here, improved alignments between the EPI and MP2RAGE scans are clearly observed in the distortion-corrected case (Fig. 6b) when compared to the original, uncorrected one (Fig. 6a); a substantial misalignment of the ventricles in the boundaries between the two scans can be found in the original EPI case. The aforementioned ventricle areas are extracted from the EPI images using signal thresholding, and the results from both the original EPI (red-yellow) and corrected EPI (blue-light blue) images are overlaid onto the MP2RAGE scan in Fig. 6c. This representation clearly illustrates the substantial improvements in co-registration achieved through distortion-correction. Specifically, in the sagittal presentation, the geometric deformations present in the frontal lobe of the original EPI image (i.e. shrunken structures) are demonstrated to be effectively reduced in the distortion-corrected case. As a result, the boundary between the cingulate gyrus and corpus callosum is also well aligned between the EPI and MP2RAGE scans, as indicated by the regions marked by red arrows.

Table 2 presents the results of the calculated LPIPS index obtained from the five-subject datasets. For all examined ROIs (GM, WM and CSF), a substantial reduction in the LPIPS index (i.e. greater similarity between the EPI and MP2RAGE scans) was observed in the distortion-corrected results when compared to original image across all subjects.

**Table 2.** Structural similarity between EPI and MP2RAGE (LPIPS index). On average, the distortion-corrected EPI showed a substantial reduction in the LPIPS index compared to the original EPI, with decreases of 7.85%, 5.03%, and 7.83% for GM, WM, and CSF, respectively.

| **Subject** | **Original EPI** |  |  | **Distortion-corrected EPI** |  |  |
| --- | --- | --- | --- | --- | --- | --- |
|  | **GM** | **WM** | **CSF** | **GM** | **WM** | **CSF** |
| **1** | 0.187 | 0.159 | 0.153 | 0.167 | 0.145 | 0.142 |
| **2** | 0.183 | 0.152 | 0.170 | 0.174 | 0.145 | 0.162 |
| **3** | 0.186 | 0.159 | 0.151 | 0.177 | 0.152 | 0.143 |
| **4** | 0.218 | 0.176 | 0.195 | 0.194 | 0.172 | 0.177 |
| **5** | 0.179 | 0.148 | 0.162 | 0.167 | 0.145 | 0.142 |
| **Average** | 0.191 | 0.159 | 0.166 | 0.176 | 0.151 | 0.153 |

### 3.4. Analysis of fMRI data

Figure 7 displays activated voxels obtained from the original (red-white) and distortion-corrected (cyan-magenta) fMRI data sets, overlaid on the respective co-registered MP2RAGE scan. Significant differences in the distribution of activated voxels are clearly visible between the original and distortion-corrected results (Fig. 7a). In separate representations of the original and distortion-corrected results (Figs. 7b and c), the distortion-corrected case demonstrates improved alignment of activated voxels within GM regions, whereas the original data show that the activated voxels are partially identified outside the GM regions. This is clearly seen in the enlarged representation of the selected rectangular ROIs (see yellow arrows). Table 3 presents the ratio of activated voxels located in the GM and non-GM regions (*r_GM_*and *r_Non-GM_*). A substantial increase in *r_GM_*was observed in the distortion-corrected case when compared to the original, consistently across all five subjects. The mean ± SD of *r_GM_*in the distortion-corrected case was 88.58 ± 2.51%, which was, on average, 8.33% higher than that of the original (80.25 ± 1.69%).

**Figure 7.**
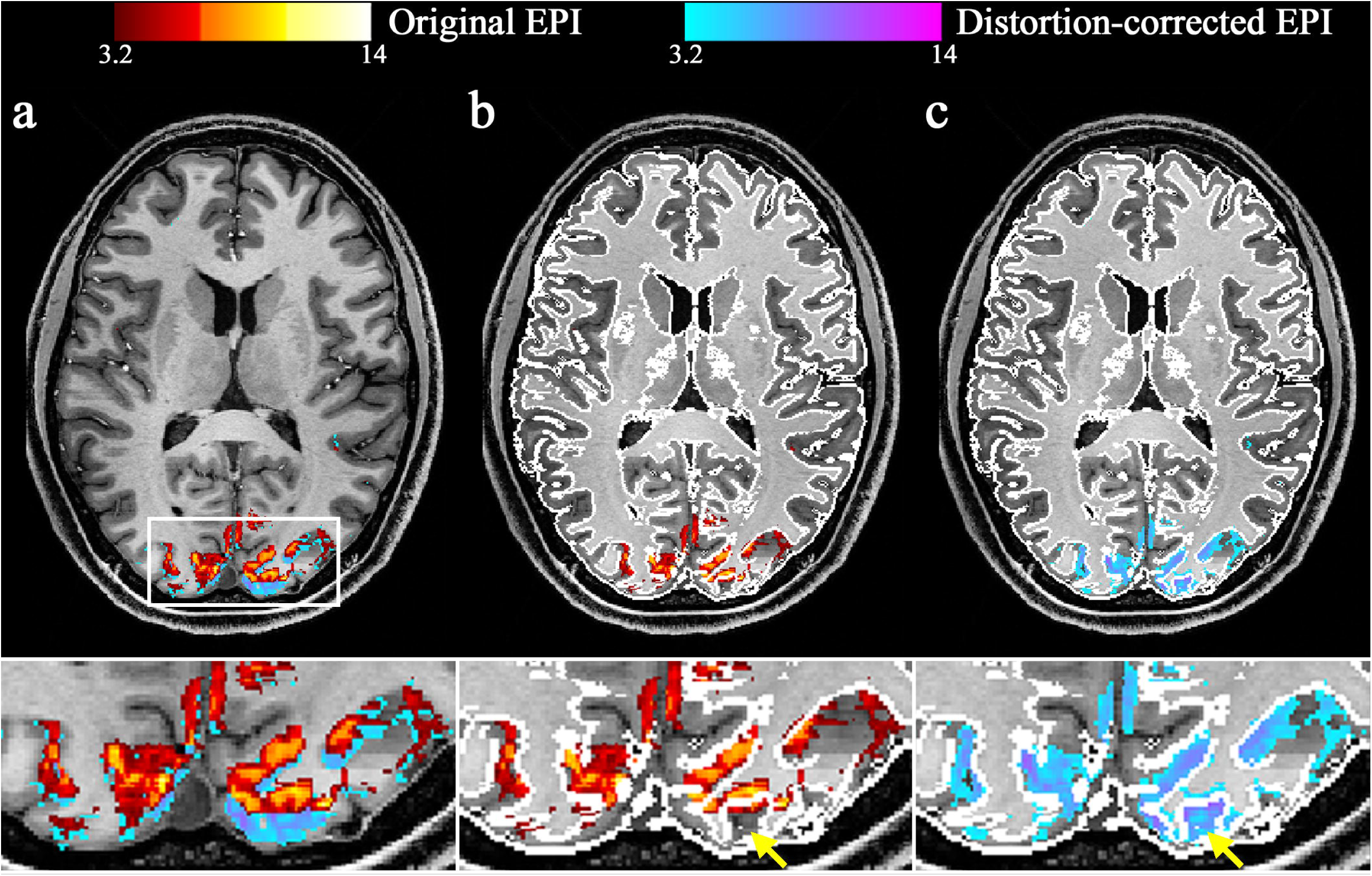
Results of first-level analysis for visual fMRI (uncorrected p < 0.001). (**a**) The original (denoted in red-white) and distortion-corrected (denoted in cyan-magenta) results are overlaid on the co-registered MP2RAGE scan, revealing substantial differences in the localisation of activated voxels at the selected slice locations. A separate representation of the (**b**) original and (**c**) distortion-corrected results with the GM contour (denoted in white) further demonstrates that the activated voxels are more precisely localised along the cortical ribbon in the distortion-corrected case; see the enlarged depiction of the selected ROIs (see yellow arrows).

**Table 3.** The ratio of activated voxels located in the GM and non-GM regions (*r_GM_* and *r_Non-GM_*). The results from distortion-corrected EPI show substantial increases in *r_GM_*(88.58 ± 2.51%) when compared to that from original, uncorrected EPI (80.25 ± 1.69%).

| Subject | Original EPI |  | Distortion-corrected EPI |  |
| --- | --- | --- | --- | --- |
| | $r_{GM} (\%)$ | $r_{Non-GM} (\%)$ | $r_{GM} (\%)$ | $r_{Non-GM} (\%)$ |
| 1 | 78.96 | 21.04 | 88.86 | 11.14 |
| 2 | 78.50 | 21.50 | 86.43 | 13.57 |
| 3 | 82.78 | 17.22 | 89.09 | 10.91 |
| 4 | 80.73 | 19.27 | 86.16 | 13.84 |
| 5 | 80.30 | 19.70 | 92.38 | 7.62 |
| mean ± SD | $80.25 \pm 1.69$ | $19.75 \pm 1.69$ | $88.58 \pm 2.51$ | $11.42 \pm 2.51$ |

Figure 8 shows the histogram distribution of t-values obtained from the original and distortion-corrected fMRI data sets, presented in blue and orange, respectively. Visual inspection of the histogram distribution suggests that the original and distortion-corrected results exhibit highly similar features across all five subjects, implying that the distortion correction had no significant impact on the overall distribution of functional activation. Table 4 presents the computed correlation coefficients (ρ*_HO,HC_*) and KS statistics (*D* and *p*-value) between the two distributions. For all subjects, ρ*_HO,HC_* exceed 0.99, and the *p*-value was greater than 0.05, with the *D* value being close to zero, indicating no statistically significant difference between the two distributions.

**Figure 8.**
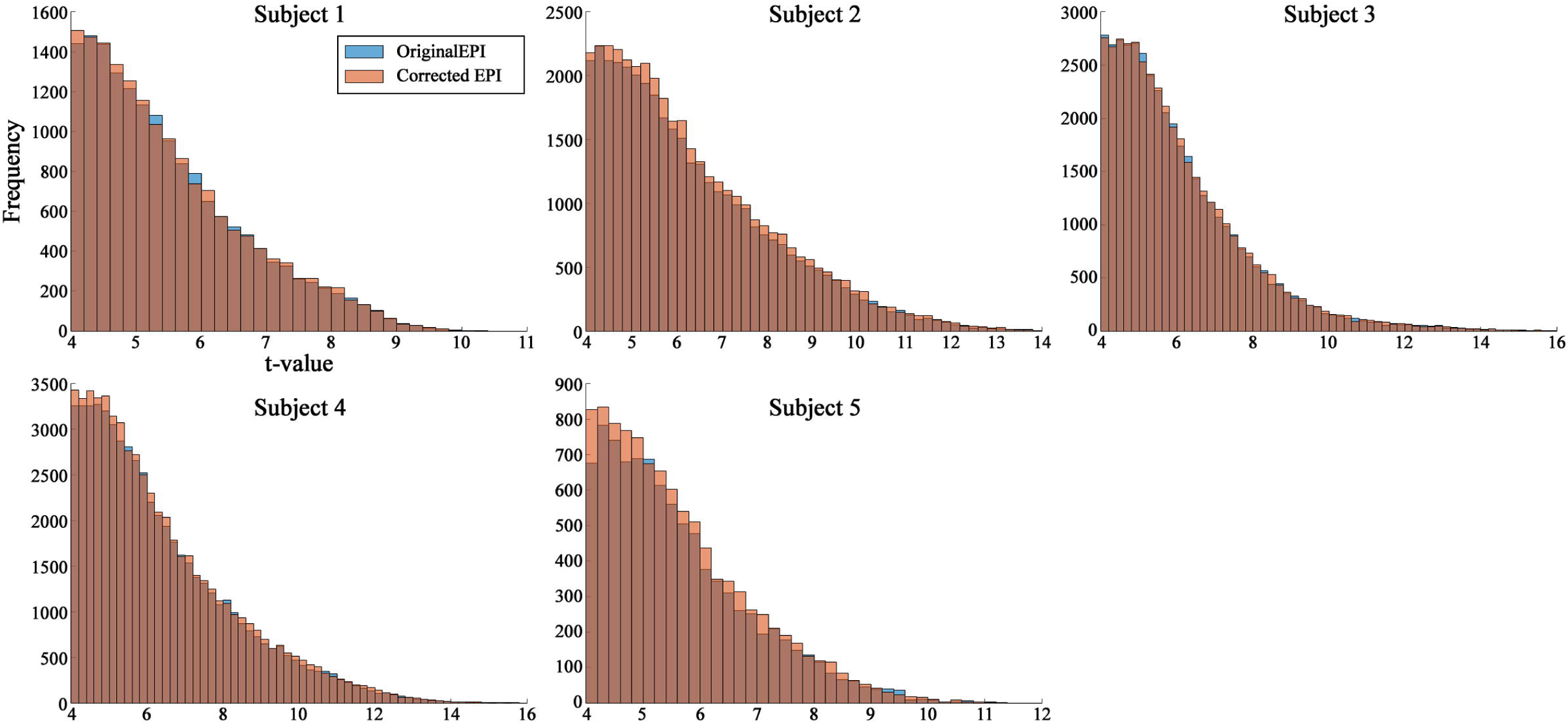
Analysis of histogram distribution from five subjects. For each subject, the histogram distribution of t-values obtained from the original (denoted in blue) and distortion-corrected (denoted in orange) fMRI data sets are depicted on the same plot, with the computed Pearson correlation coefficient. The histogram distribution and correlation values (ρ > 0.99 for all subjects) show a high degree of similarity, implying that the impact of distortion correction on the overall distribution of functional activation was not significant.

**Table 4.** Results of statistical measures used to evaluate the difference between the two functional distributions before and after distortion correction.

| Subject | Correlation coefficient ($\rho_{HO,HC}$) | KS test ($D$ and $p$-value) | |
| --- | --- | --- | --- |
| 1 | 0.9990 | 0.0079 | 0.6819 |
| 2 | 0.9992 | 0.0094 | 0.0687 |
| 3 | 0.9996 | 0.0075 | 0.1941 |
| 4 | 0.9994 | 0.0059 | 0.2637 |
| 5 | 0.9996 | 0.0090 | 0.8211 |

## 4. Discussion

This work proposes an EPI scheme that acquires both original and reversed PE-direction EPI in a single fMRI session and demonstrates its applicability for the distortion correction of submillimetre fMRI data at 7T. When compared to the standard scheme, the proposed approach effectively reduces additional acquisition time, RF energy accumulation, and subject motion accrued during the acquisition between the two different PE directions. This advantage becomes more prominent when the parallel imaging calibration scan is acquired using the SPGR sequence, a method commonly adopted at ultra-high fields for more robust calibration of the reconstruction kernel. In this study, the SPGR sequence was configured with 72 ACS lines, a relatively high number intended to enhance reconstructed image quality. Albeit at the cost of an increase in the scanning time, this approach serves as a straightforward solution unless more elaborated sampling methods (Park et al., 2005; Wang et al., 2014) or advanced kernel calibration techniques (Aja-Fernández et al., 2015; Sheng et al., 2021; Chang et al., 2022) are implemented. In this regime, the advantages of the proposed EPI become more pronounced, as it eliminates the need for repeated acquisition of the calibration scan, which would otherwise significantly prolong the acquisition in the standard scheme.

The geometric distortion in the original fMRI data was significantly reduced through the blip-up-blip-down method. While other distortion-correction techniques, such as the B_0_-field map or PSF methods, have been previously demonstrated, a comparative assessment of various distortion-correction methods is not the focus of the current work, as this topic has already been explored in prior research (Zeng and Constable 2002; Wang et al., 2017; Patzig et al., 2021; Malekian et al., 2023). This study primarily focuses on a thorough evaluation of distortion correction performance in the context of submillimetre resolution and its potential impact on fMRI analysis. To achieve this, various qualitative and quantitative evaluation criteria were proposed, including assessments of spatial resolution (σ_deg_), co-registration accuracy (LPIPS index), functional mapping accuracy (*r_GM_*), and functional distribution (ρ_HO,HC_, *D*, and *p*-value).

One frequently addressed concern associated with distortion correction is the potential loss of spatial resolution arising from the unwarping process, which involves resampling or repositioning of voxels (Renvall et al., 2016; Kashyap et al., 2018). However, the analysis results demonstrate that the potential degradation in spatial resolution caused by the distortion-correction process was minimal, as confirmed by data from five subjects. This work employed ANTs to implement the blip-up-blip-down distortion correction. In the community, other software packages are also available for the same purpose, including 3DQwarp (AFNI) (https://afni.nimh.nih.gov), COPE (Brain Voyager) (https://www.brainvoyager.com), and TOPUP (FSL) (https://fsl.fmrib.ox.ac.uk/fsl), all of which have been employed in numerous previous submillimetre fMRI studies. In future work, the performance of these alternative distortion-correction software will be assessed in comparison to the ANTs, with the aim of determining the most optimal method for high-resolution fMRI.

Accurate mapping of detected neural signals onto the correct anatomical references requires precise co-registration between functional and anatomical scans, as measured by EPI and MP2RAGE. In particular, submillimetre fMRI, which allows for the characterisation of cortical depth-dependent functional activities, demands a clear definition of cortical regions, typically achieved using the structural information derived from the anatomical scan (Margalit et al., 2020; Jia et al., 2021; Deshpande et al., 2022). The distortion correction demonstrated here exhibited significant improvements in the co-registration between the EPI and MP2RAGE scans, as confirmed by both visual inspection as well as the quantitative

LPIPS index. However, the distortion-corrected EPI images still suffer from the susceptibility-induced signal loss, which can be seen in regions around the frontal lobe. This can be reduced by employing dedicated RF pulses, also known as tailored RF methods (Yip et al., 2006; Zheng et al., 2013), or utilising an EPI-readout shortening technique such as EPI with Keyhole (EPIK) (Shah and Zilles, 2003, 2004; Zaitsev et al., 2001), the use of which has been demonstrated in a number of fMRI studies (Yun et al., 2013, 2019; Yun and Shah, 2017) including those focused on delineating the cortical depth-dependent profiles (Yun et al., 2022; Pais-Roldán et al., 2023). In this study, the target functional area was the visual cortex, where the susceptibility-induced signal loss is expected to be minimal. This allowed the investigation of the effect of distortion correction on functional activation with little concern for the signal loss issue.

The improved co-registration directly contributed to enhancing the accuracy of functional mapping onto the GM. Here, the distortion correction did not significantly affect the distribution of functional activation, as verified by the histogram distribution and the statistical assessment using the KS test. In the distortion-corrected case, a significant increase in *r_GM_* was observed when compared to that from the original, uncorrected data. Nevertheless, a fraction of the activated voxels still appeared in the non-GM regions, with an average *r_Non-GM_* of 11.42% across the five-subject data sets. Several possible factors contribute to the presence of non-GM activated voxels. One explanation is imperfect distortion correction and the subsequent errors in co-registration. Another potential cause is false positive activation induced by motion or physiological fluctuations, such as respiration or cardiac pulse (Birn et al., 2006; Bright et al., 2013). The impact of physiological noise can be controlled to some extent by simultaneously recording the physiological signals during fMRI and regressing them out from the fMRI time-series data (Birn et al., 2006). Another factor may be the superficial bias effect typically seen in the gradient-echo BOLD methods. Various approaches have been proposed to reduce this effect, including the use of prior information on venous structures (Koopmans et al., 2010; Kay et al., 2019) or phase-data-based regression methods (Menon et al., 2002; Stanley et al., 2021). We anticipate that when our fMRI setting is further combined with the physiological-noise or the superficial-bias control methods, an even higher *r_GM_* will be achieved.

A distortion-matched anatomical scan, the readout of which is performed using the same EPI sequence as for fMRI, can be used as an alternative to the distortion correction method (Chai et al., 2021; van der Zwaag et al., 2018). This anatomical scan exhibits distortions identical to the fMRI scan, ensuring errorless co-registration between the two scans. This approach makes the co-registration process more straightforward compared to the scenario involving two different sequence modalities (e.g. EPI and MP2RAGE) (Kashyap et al., 2018; Navarro et al., 2021). However, due to the distorted anatomical reference, its application can be limited for the analyses that rely on accurate anatomical information, such as the morphometric measurement of cortical thickness or the cortical surface-based atlasing (Polimeni et al., 2018, Yun et al., 2024). The distortion in this anatomical scan can also be corrected using the reversed PE method. In this case, the same simultaneous acquisition scheme proposed in this work can be effectively deployed to minimise the total acquisition time and energy accumulation.

One limitation of the reversed PE method is its relatively time-consuming process. In our fMRI data, which consists of 96 temporal volumes, each containing 9.7 M voxels (i.e. 288 × 288 × 117 slices), the processing time on a standalone PC (Intel Core i7-8850H CPU 2.60 GHz with 32GB RAM) was approximately 6.5 hours. The processor used here has a single socket and 12 parallel processing threads, which were fully exploited during the ANTs computation. However, given that our processor is relatively old (Intel Core 8th generation, released in Apr. 2018), the use of a more powerful processor with multiple sockets under a workstation environment (e.g. eight sockets, 48 threads with Intel Xeon Processor E7, yielding 384 parallel processing threads), could drastically reduce the processing time, allowing for a timely reconstruction for entire submillimetre fMRI data with distortion correction.

In conclusion, this work presents a novel EPI scheme that simultaneously acquires reversed PE data in a single measurement, and its utility for submillimetre (0.73 × 0.73 mm^2^) fMRI has been demonstrated at 7T. This strategy effectively spares another acquisition time and RF energy accumulation for the parallel imaging calibration scans, making the method particularly applicable for time-critical clinical applications. The performance of distortion correction was evaluated in direct comparison with original, uncorrected data using the various quantitative and qualitative assessment criteria proposed here. The results demonstrate improved functional mapping accuracy without significant degradation of spatial resolution or alternation of the functional activation distribution. This provides direct insight into the effectiveness of distortion correction for submillimetre fMRI, as demonstrated in this study.

## Acknowledgement

The authors would like to thank the volunteers for their participation in this work and Claire Rick for English proofreading.

## Declaration of generative AI in scientific writing

During the preparation of this work, the author used ChatGPT-3.5 (free version) solely to enhance the readability and language of the manuscript. The initial text was carefully written by the authors, and after receiving ChatGPT’s suggestions, the author reviewed and edited the content as needed. No paragraphs were originally generated by AI. The authors take full responsibility for the content of the published article

## Author Contributions

S. Y. designed the research; S.Y. developed the proposed MR imaging sequence and the corresponding image reconstruction software; S.Y. performed data acquisition and analysis; S.Y. wrote the draft of the manuscript and revised it; E.G. introduced the geometric distortion using the ANTs software, and reviewed the manuscript. J. L. performed data acquisition and analysis, and reviewed the manuscript. N.J.S. and S.Y held critical discussions; N.J.S. reviewed and revised the manuscript; N.J.S supported the maintenance and use of the 7T MRI scanner.

