## Supplementary File for "The Simultaneous Measurement of Reversed Phase-Encoding EPI in a Single fMRI Session: Evaluation of Geometric Distortion Correction in Submillimetre fMRI at 7T"

**Supplementary Figure**

**1.1 Entire reconstructed slices of distortion-corrected EPI**

The figure below shows distortion-corrected results from a single-volume functional scan for the entire axial slices (117 slices) acquired with our submillimetre protocol (0.73 × 0.73 mm2). All slices were well reconstructed without any significant visible loss of spatial resolution, demonstrating reliable distortion correction performance across all slice locations.

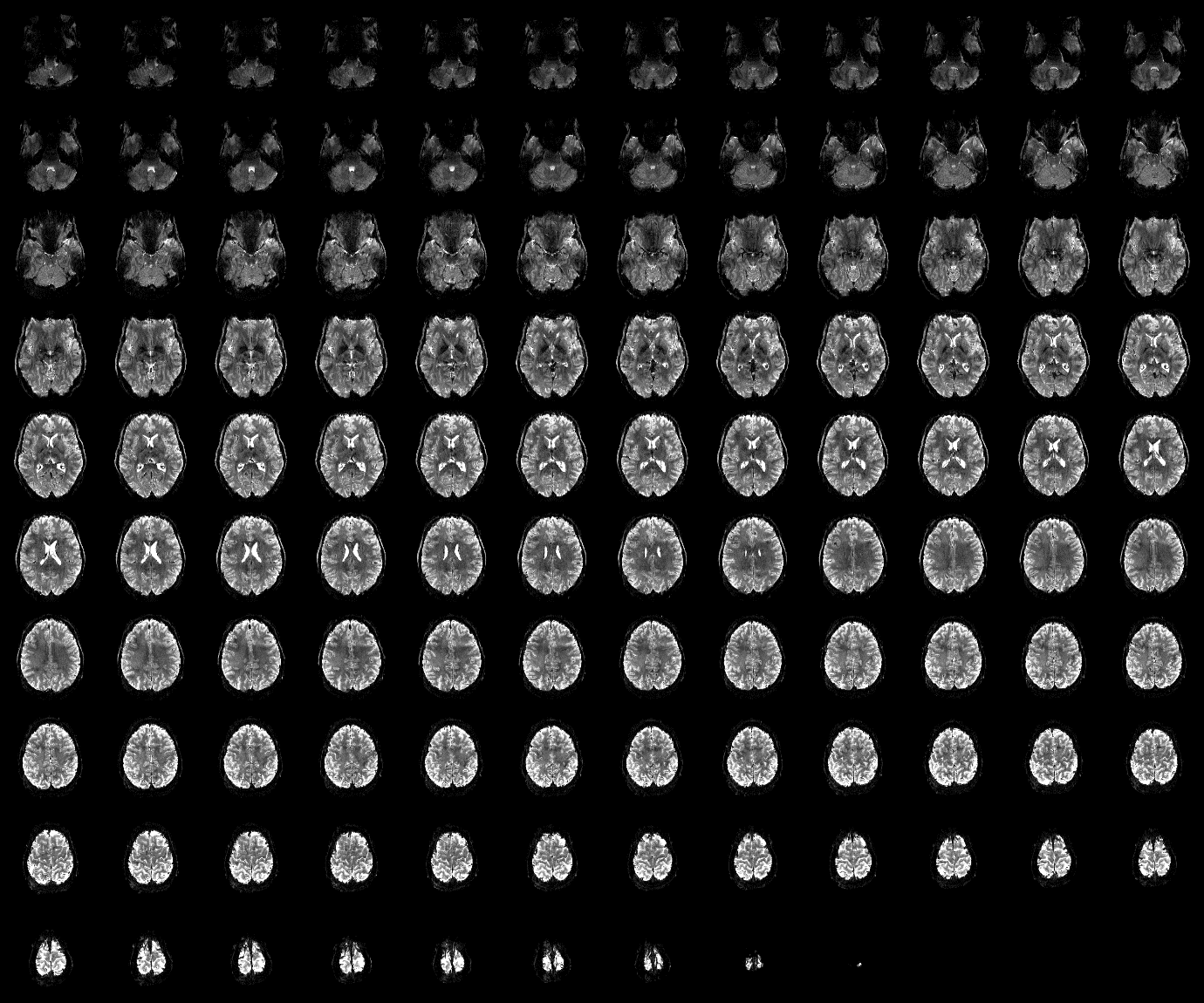

**Supplementary Figure 1**. Distortion-corrected EPI of the submillimetre protocol (0.73 × 0.73 mm2) for the entire 117 axial slices (1 mm thickness).

**1.2 Acquisition time and SAR of the proposed scheme**

The proposed scheme enables the reconstruction of both original and reversed-PE EPI images using the same calibration data acquired before EPI acquisition, offering a more optimised configuration in terms of the total acquisition time and SAR, when compared to the standard scheme. For quantitative evaluation, the reduction in acquisition time and SAR was simulated using the Siemens sequence development platform, ‘IDEA’ (Integrated Development Environment for Applications), under the baseline version of N4_VE12U_LATEST_20181126.

Supplementary Table 1 shows the acquisition time and SAR required for the parallel imaging calibration scan when acquired using the SPGR sequence and compares them with those needed for the entire fMRI acquisition. The results were obtained for a range of ACS lines (i.e. 24, 48, and 72) and TRs (i.e. 10, 15, and 20 ms) from the SPGR sequence. It is important to note that the SAR values shown in the table represent the time-averaged radio frequency (RF) power (i.e. W), calculated at a commonly used flip angle of 15°. The table reveals that the required acquisition time increases with higher numbers of ACS lines or longer TR values. While the ACS acquisition time constituted a relatively small portion compared to the total acquisition time, it reached as high as 192 s when acquired with a TR of 20 ms and 72 ACS lines. The implications of using a relatively long TR and a large number of ACS lines will be discussed later.

Furthermore, the table demonstrates that SAR decreases for longer TR values, as readily expected. However, SAR also decreases for a greater number of ACS lines. This is due to the fact that the RF energy per unit time dissipated in the parallel imaging calibration scan is much smaller than that of the main fMRI scan, as it is implemented with large flip angles for the main excitation (90°) and the fat-saturation module (110°). Conversely, when the calibration data are acquired with a relatively short TR and small ACS lines (i.e. 10 ms with 24 ACS lines), SAR accounts for as much as 46% of the total SAR required. The above results suggest that the proposed simultaneous acquisition scheme can significantly reduce the redundant SAR and acquisition time for the acquisition of reversed PE EPI.

**Supplementary Table 1**. Acquisition time (s) and time-averaged power (W) required for the parallel imaging calibration scan. The results are shown for a range of ACS lines (24, 48, and 72) and TR (10, 15, and 20 ms), when a SPGR sequence is implemented with a flip angle of 15°. For each case, the time and power required for the entire measurement are also provided, along with the proportion occupied by those for the parallel imaging calibration scan (see the percentages in the parentheses).

| **ACS lines** | **TR (ms)** | | | | | |
| --- | --- | --- | --- | --- | --- | --- |
|  | **10** | | **15** | | **20** | |
| **24** | 39.78 / 421.29 s  2.51 / 5.45 W | (9.44 %)  (46.10 %) | 59.67 / 441.18 s  2.01 / 5.21 W | (13.53 %)  (38.50 %) | 79.56 / 461.07 s  1.67 / 4.98 W | (17.26 %)  (33.46 %) |
| **48** | 67.86 / 449.37 s  1.93 / 5.13 W | (15.10 %)  (37.62 %) | 101.79 / 483.30 s  1.46 / 4.77 W | (21.06 %)  (30.67 %) | 135.72 / 517.23 s  1.18 / 4.46 W | (26.24 %)  (26.43 %) |
| **72** | 95.94 / 477.45 s  1.59 / 4.85 W | (20.09 %)  (32.81 %) | 143.91 / 525.42 s  1.17 / 4.41 W | (27.39 %)  (26.61 %) | 191.88 / 573.39 s  0.93 / 4.04 W | (33.46 %)  (22.99 %) |
